# Distinct mechanisms of germ cell factor regulation for an inductive germ cell fate

**DOI:** 10.1101/2022.02.04.479164

**Authors:** Stephany Foster, Nathalie Oulhen, Tara Fresques, Hossam Zaki, Gary Wessel

## Abstract

Specification of primordial germ cells (PGCs), the lineage which gives rise to eggs and sperm, is essential for sexually reproducing organisms. The mechanism by which animals specify their PGCs generally falls into two categories: inherited or inductive. The inductive mechanism, used by mammals, relies on cell signaling interactions to direct a subset of embryonic cells to a germ cell fate. Previous work suggested that sea star embryos, which develop in simple culture and are markedly transparent, also use inductive mechanisms to specify their germline. The germ cell factors Nanos and Vasa become restricted during early development into a localized region of cells within the posterior enterocoel (PE), the presumptive germline. Nodal signaling was observed to negatively regulate Vasa and Nanos mRNAs outside of the PE and restrict the germline to the PE. Here we employed single cell RNA sequencing to identify the transcriptional program of germ cells and their changes during development. We never see Nodal pathway members within Nanos/Vasa positive cells in the region known to give rise to the PE, and instead see members of the Wnt-signaling pathway and the FoxY family of transcription factors. We learned that Wnt and Delta/Notch signaling enhances expression of both Nanos and Vasa, whereas a test of cell interactions reveals that Nanos and Vasa are regulated distinctly. This work provides insights into the sequence of events that leads to PGC specification and enables deeper mechanistic studies in a tractable *in vivo* model.

**Highlights:**

1. Single-cell RNA-sequencing of sea star embryos demonstrates temporal differences in cell fate commitment among echinoderms.
2. Sea urchin and sea star embryos appear to ascribe their germ line by two extreme different mechanisms but share similar pathways in regulation of the germline genes.
3. Expression of the germline factors, Vasa and Nanos, is regulated by distinct mechanisms in the sea star.
4. Germline induction in the sea star uses similar signaling mechanisms as mammals.

## Introduction

Sexually reproducing organisms have a distinct germline, the cell lineage that will give rise to either eggs or sperm in the adult, enabling a continuation of species. Two major mechanistic distinctions have been identified for specification of the primordial germ cells, the cells committed early in development to the germline. These are referred to as the inductive and the inherited mechanisms also referred to as epigenesis and preformation, respectively [1]. In the inherited mechanism, used by species including the insect *Drosophila melanogaster*, the frog *Xenopus laevis*, and zebrafish *Danio rerio*, localized maternal RNA transcripts and proteins in the egg are inherited by a particular set of cells in the embryo, which then give rise to gametes in the adult [1, 2]. In contrast, in the inductive mechanism, the germline is formed later in development through zones of cell-cell signaling. Current evidence suggests that a combination of activating and inhibitory signals converges to induce germline specification in a restricted region of the embryo. This mechanism is used by species including mammals, the cricket *Gryllus bimaculatus*, and the amphibian *Ambystoma mexicanum* (axolotl). Studies in organisms across many different phyla place the inductive mechanism as the ancestral mechanism of germline formation [1]. However, the full network of cell signaling pathways and their targets which direct the ancestral mode of germline formation remains poorly understood. A role for BMP signaling in germline specification is thought to be conserved. Loss of BMP signaling results in loss of primordial germ cells in mouse and cricket embryos [3, 4]. Wnt signaling was identified to induce PGC specification in both mouse and axolotl, although the specific Wnt ligand is not conserved [3, 5, 6].

The sea star, *Patiria miniata*, follows an inductive mechanism of germline formation [7]. This is in contrast to how its close relative, the sea urchin, forms its germline which instead relies on maternal factors. This divergence provides an important comparison between organisms within a taxon. Further, the sea star provides a transparent, tractable embryo, closely related to chordates, for *in vivo* analysis. Expression of the highly conserved germline marker genes Nanos and Vasa, among others, has been characterized [8]. Nanos and Vasa expression is seen in a vegetal ring at mid-gastrula stage. Later, Nanos and Vasa expression is lost in the ventral gut and then the right side of the developing gut at late gastrula stage. Finally, at the larval stage, Nanos and Vasa expression is restricted to the dorsal left side of the gut and those cells form the posterior enterocoel (PE). We refer here to the PE as the presumptive germline as previous work has shown that removal of the PE results in significantly fewer germ cells in larva compared to controls [9]. The PE also accumulates Nanos and Vasa transcripts selectively in the larva supporting its germline contention. The lineage of cells in the PE though, have not yet been traced through development to adulthood.

Initially, Nanos and Vasa RNA are expressed broadly in the forming gut, and Nodal signaling inhibits expression of the germline genes, restricting germline gene expression eventually to the left side. A Nodal signaling gradient on the ventral side at blastula stage, and on the right side in late gastrula stage are responsible for restriction of Nanos and Vasa to the dorsal left side, in the PE [7]. While Nodal has been identified to play a role in restricting the germline by inhibiting expression of the germline genes, the signal or signals that activate germline gene expression is unknown.

Single-cell RNA-sequencing (scRNA-seq) is a powerful tool used to identify individual cells that make up an entire embryo. Single-cell RNA sequencing has identified germline cells of *Xenopus*, zebrafish, and sea urchin embryos, and has been employed to identify transcriptomic changes in germline cells with development in male and female mice [10–14]. We performed scRNA-seq in the early sea star to identify the cell states expressing germline factors in the embryo, to test the cell signaling pathway components responsible for their fate, and to leverage analysis of the distinct mechanism used for germ cell formation in the sister taxon of sea urchins.

## Results

### Identification of cell states across early sea star development

Sea star embryos were cultured to six different developmental stages, 8 hpf, 10 hpf, 14 hpf, blastula, early gastrula, and mid-gastrula stage, at which point they were dissociated and processed for scRNAseq via Drop-seq [15]. All six datasets were analyzed using Seurat and integrated using Harmony, a single-cell RNA-sequencing data integration tool, to identify conserved cell types across all datasets [16, 17]. This analysis of 25,703 total cells yields 20 clusters, or cell states, across all timepoints identified by known marker gene expression (Figures 1A and 1B; Table S6). Here we use cell state to refer to a select accumulation of transcripts present in a cell as it reaches a developmental fate. In analyzing the six datasets together, it is evident that early (8-14h) and late (B-MG) cell types are transcriptionally distinct. Cell clusters of early stages (Clusters 3 and 7) decrease with development; these cells presumably are “lost” to differentiation (Figure S1). Clusters 3, 7, and 9, which make up the bulk of the earlier stage datasets, showed enrichment of CyclinA and lack of marker gene expression (Figure 1B).

**Figure 1.**
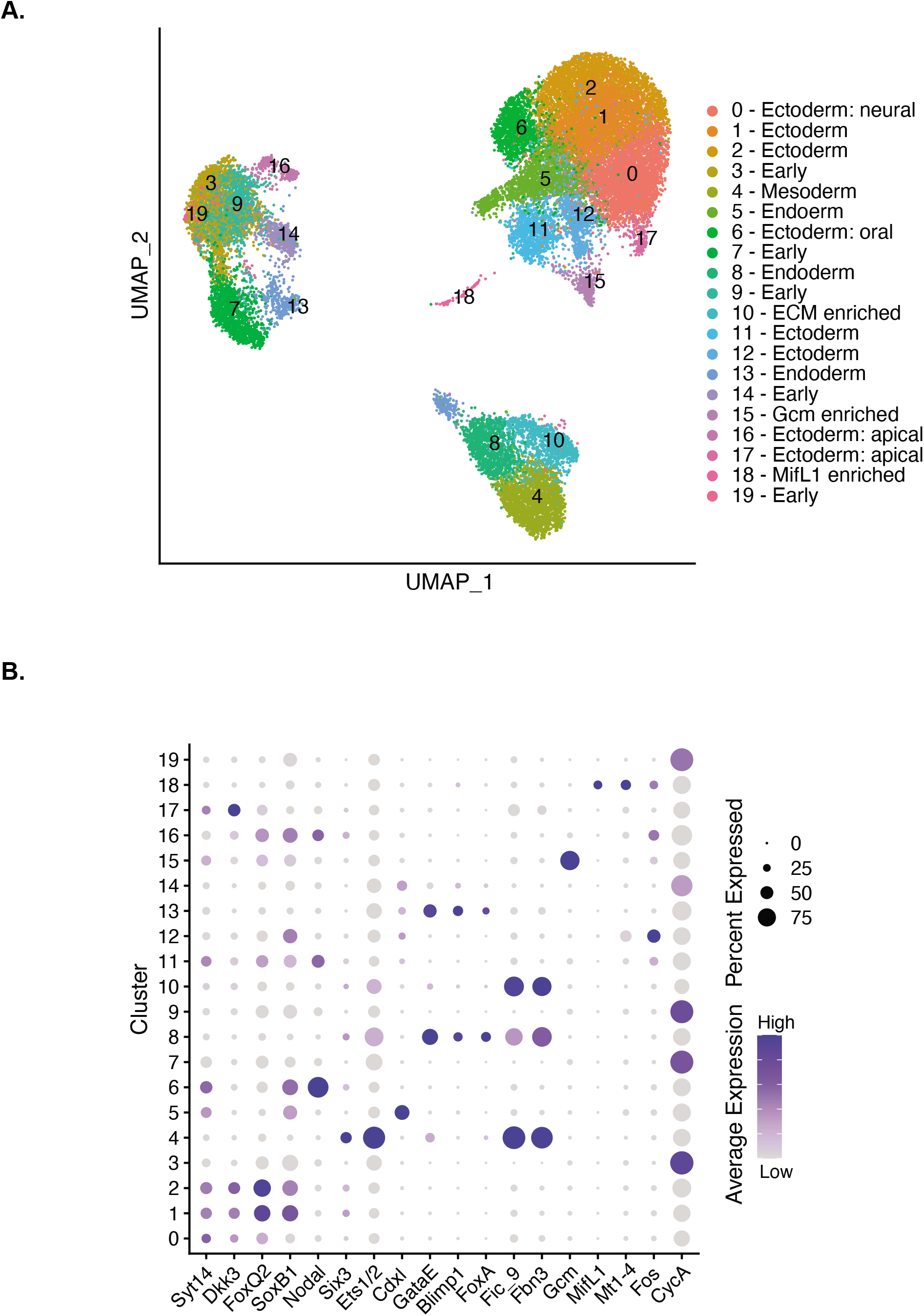
Cell states identified across early sea star development. (A) UMAP visualization of 25,703 cells after integration of six datasets. Cells colored by cell state identity based on marker gene expression. (B) Dotplot showing marker gene expression across clusters. Average gene expression level displayed by color intensity. Percent of cells expressing the marker gene conveyed by circle size.

### Blastula

We assessed the blastula and gastrula stage datasets in greater detail as gene expression in these developmental stages have been characterized previously and germline marker gene expression is present. In blastulae, we detect 10 cell states which are identified by expression of such known marker genes (Figures 2A and 2B; Tables S1 and S3). We identified ectodermal cells by expression of SoxB1 [18]. Foxq2 and DKK3 are expressed in the animal pole domain where the apical organ of the nervous system will form [19]. Cluster 0 expresses high levels of SoxB1, FoxQ2, and Dkk3. Cluster 1 expresses FoxQ2 and Dkk3 which we identify as the animal pole domain (Figure S2). While these apical ectodermal markers are widespread in the dataset, the vegetal markers are restricted to a subset of cell states. These vegetal markers include both mesodermal and endodermal progenitors. Ets1/2, a marker of the presumptive mesoderm, is expressed in the vegetal plate [20]. GataE, FoxA, brachyury, and Blimp1, all genes involved in endodermal development, will mark different regions of the embryonic gut. However, in blastulae, they are all expressed in the vegetal ectoderm, surrounding the blastopore [8, 21]. Cdxl, another endodermal marker, is also expressed as a ring, vegetally in blastulae; it is later restricted to the blastopore region in mid-gastrulae [22]. GataE, FoxA, and Blimp1 are expressed in the same cluster as Ets1/2 (cluster 2) while brachyury and Cdxl are markers of cluster 3 (Figures 2B and S2). At blastula stage, the transcription factor glial cells missing (gcm), which marks the pigment cells in a related species, the purple sea urchin shows no localized expression; gcm-positive cells are present throughout the ectoderm [23, 24]. In our dataset, cells with highest gcm expression clustered into a unique population in cluster 8 with some gcm-positive cells present within the ectodermal clusters (Figures 2B and S2).

**Figure 2.**
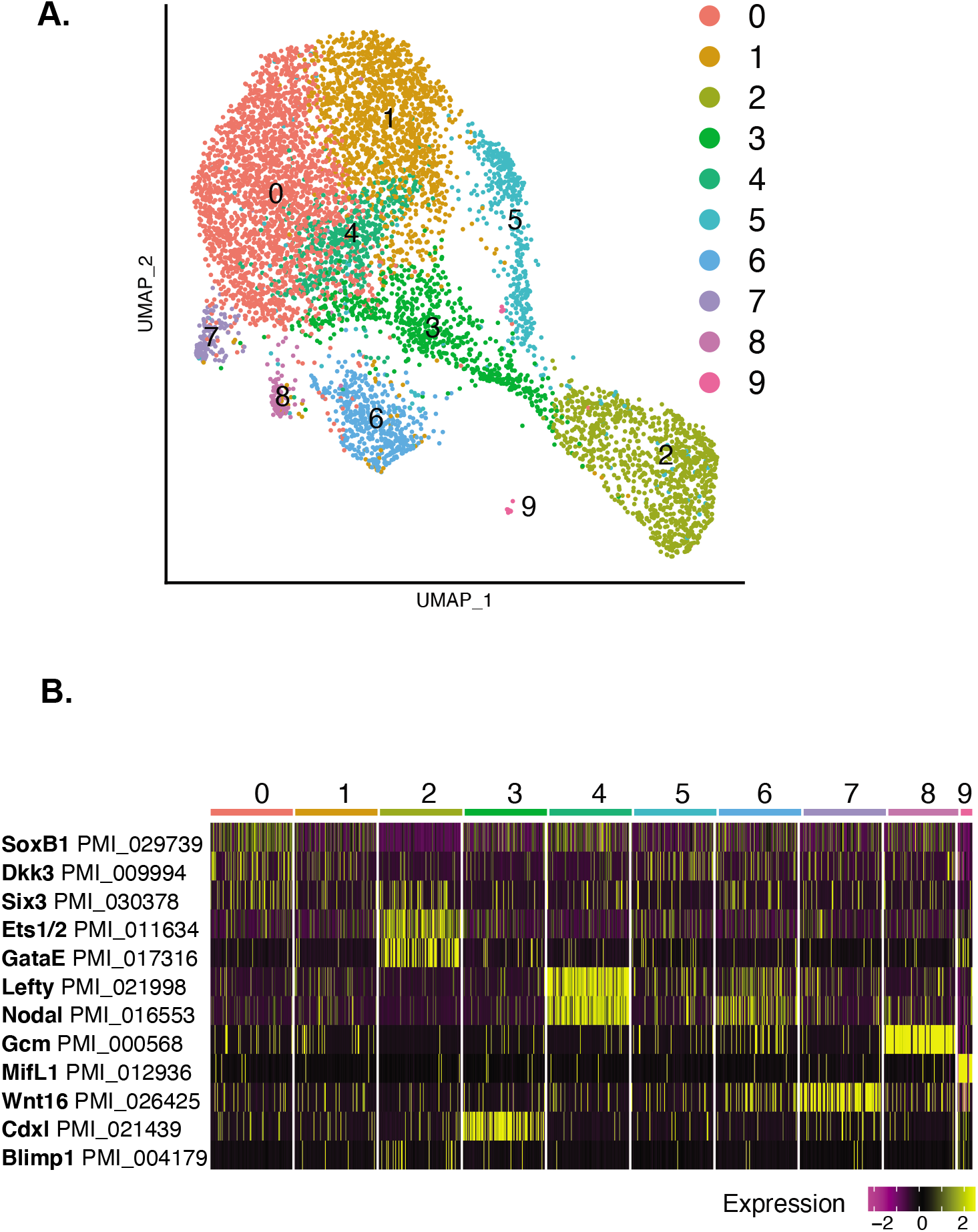
Gene expression at blastula stage. (A) UMAP plot of 7,272 blastula stage cells. Clusters colored by cell state identified by marker gene expression. (B) Heatmap of blastula stage marker gene expression.

Expression of the Wnt signaling ligands and receptors has previously been characterized in blastula and gastrula stages [25]. As signaling plays an important role in development and inductive germline specification, we sought to characterize Wnt ligand expression in our data. Our dataset showed coexpression of ligands Wnt3, Wnt16, and the Wnt receptor, Frizzled1 (Frizz), in cluster 2 (Figure S4). These three are known to be expressed in the vegetal pole at blastula stage. The ligand Wnt8 is also expressed in the vegetal pole, not overlapping with Frizz expression. Additionally, the Wnt8 expression pattern showed overlap with Wnt3 (cluster 3) and non-overlap with Wnt3 (cluster 7) mirroring what was previously reported (Figure S4) [25]. Vasa mRNA is expressed in the vegetal plate of blastulae and is particularly enriched in cluster 2, like the Wnt3 and Wnt16 ligands, and the receptor Frizz (Figure S6H) [8]. Nanos accumulation is seen by in situ hybridization later, in mid-gastrulae, but is not detected in blastulae [26]. Indeed, Nanos expression was detected at low level at blastula stage in our dataset (Figure S6A).

### Mid-gastrula

Analysis of the mid-gastrula stage dataset reveals 11 cell states (Figure 3A; Tables S1 and S4). At this stage, many characteristics of the animal ectoderm remain the same. Ectodermal cell types (cluster 0,1,2,4) are still distinguished by expression of SoxB1, FoxQ2, and Dkk3 (Figure S3). Now, expression of several vegetal markers detected at blastula stage, changes from overlapping to marking either vegetal ectoderm or different regions of the gut. In blastula stage, both Ets1/2 and GataE were expressed in the same cluster (cluster 2), at gastrula stage, Ets1/2 is particularly enriched in cluster 3, and GataE marks a different cluster (cluster 6) (Figure 2B). Ets1/2 at gastrula stage is present at the tip of the archenteron where it marks presumptive mesoderm and ingressing mesenchyme cells [20, 27]. There is some low level GataE expression in the archenteron, however, GataE is enriched in the mid- to hindgut region in mid-gastrula [21]. FoxA, which is expressed in the mid/hindgut region, is enriched in the same cluster as GataE [28]. In mid-gastrulae, brachyury expression surrounds the blastopore but is not within the blastopore [28]. Cdxl expression is restricted to the region around the blastopore [22]. Brachyury and Cdxl are both enriched in cluster 8 (Figure S3).

**Figure 3.**
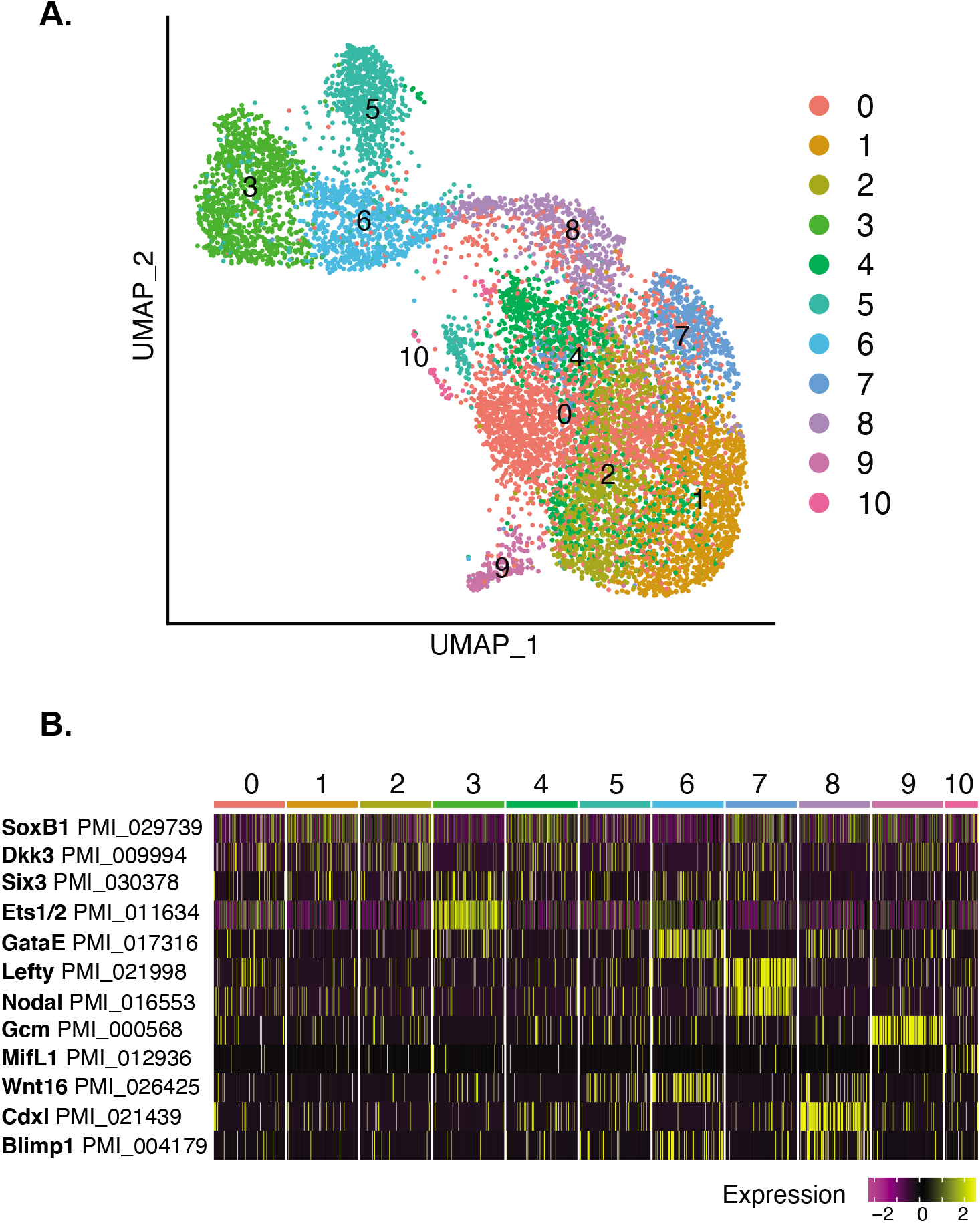
Gene expression at mid-gastrula stage. (A) UMAP plot of 10,448 mid-gastrula stage cells. Clusters colored by cell state identified by marker gene expression. (B) Heatmap of mid-gastrula stage marker gene expression.

Lack of unique marker gene expression makes cluster 5 difficult to identify. Cluster 5 showed enrichment of ECM protein expression, including fibrinogen (PMI_003448 Fic_9) and fibrillin (PMI_026729 Fbn3) (Table S4). Similar to blastula stage, one cluster is marked by high gcm expression, cluster 9. A small number of cells cluster on their own (cluster 10) and these cells are marked by MifL1, macrophage inhibitory factor like 1 (Figure 3B). MIF plays a role in immune function in vertebrates [29, 30]. Coexpression of gcm and another MIF, mif5, was previously reported in sea urchin larvae [31]. Cluster 10 also expresses Mt1-4, a metalloprotease, and Fos, a transcription factor, both enriched in the sea urchin primary mesenchyme cells (PMCs) (Figure S5) [32, 33]. Further studies are necessary to reveal what role these genes play in sea star development as sea star embryos lack PMCs and pigment cells.

The sea star embryo is known to have left/right morphological asymmetry by gastrula stage. A Nodal signaling gradient on the right side of the embryo is responsible for restricting PE formation to the left side [7]. Nodal expression, along with Lefty, is enriched in one cluster (cluster 7) which we identify as the right-side ectoderm (Figure 2B). In contrast to Nodal signaling, Wnt pathway component expression is present in the same region as Nanos and Vasa RNA expression in the early embryo and in the forming gut during the gastrula stage. In situ hybridization shows enrichment of Nanos and Vasa transcripts at the top of the archenteron and in a vegetal ring in the hindgut region at mid-gastrula stage [7]. Six Wnt ligand proteins and three proteins for Frizzled (Fz), the Wnt receptor, are present in the *P. miniata* transcriptome [25]. Among the six Wnt ligands, RNA expression patterns of Wnt3, Wnt8, and WntA/4 show overlap with the region of Nanos and Vasa RNA expression in the gastrula stage embryo [7, 25]. Among the three Wnt receptors, Fz1/2/7 RNA expression is concentrated in the forming gut of the early gastrula [25]. The three signaling ligands, Wnt3, Wnt8, and WntA/4, show appropriate expression patterns to be candidates for initiating germline specification by interacting with the Fz1/2/7 receptor, due to their tight colocalization with the Nanos and Vasa RNA expression pattern in the early embryo. At mid-gastrula, the Wnt signaling ligands have been described as having a nested gene expression pattern. Wnt16 shows most vegetal expression followed by Wnt3 and then Wnt8 along the ectoderm. In our dataset, Wnt16 enrichment is seen in cluster 6 with Wnt3 expression. Wnt3 is also coexpressed with Wnt8 in cluster 8. Frizz is present in cluster 3, which we identify as the tip of the archenteron (Figure S4) [25].

### Derivation of the germline cells

We focused our analysis on the mid-gastrula stage, where Nanos and Vasa are both expressed and will be restricted to the region of PE formation. Nanos expression is detected in 267 total cells, co-expression of Nanos and Vasa is seen in 212 cells. Nanos expression is much less abundant compared to Vasa; Vasa expression is seen in all clusters at mid-gastrula stage but expression in the hindgut cluster (cluster 8) was of greatest interest to us (Figure S6H). We took a closer look at the Vasa-positive cells in the hindgut cluster and compared gene expression in the hindgut cells that express Vasa to those that do not. In extracting a list of differentially expressed genes in Vasa-positive cells of the hindgut cluster, we find the transcription factor FoxY3 (Table S5). In situ hybridization analysis shows FoxY3 is expressed in the same regions as both Nanos and Vasa at mid-gastrula stage (Figure S7A). FoxY3 expression is enriched in clusters 3, 6, and 8, which correspond to the archenteron and gut region, similar to Vasa expression (Figures S6H and S6I). A role for FoxY was also previously identified in regulating Nanos expression downstream of Delta/Notch signaling in the sea urchin [34]. ScRNA-seq analysis reveals that Vasa and FoxY3 are co-expressed (Figure S7B). Wnt8 and Wnt3 are expressed in the cells neighboring the Vasa, FoxY3-positive cells; Wnt3 is also expressed in the Vasa, FoxY3-positive cells, along with Wnt16 and Wnt1 (Figure S7C; Table S5). Moreover, at this stage, Vasa-positive cells express hindgut identity genes such as GataE and FoxA, and the homeobox gene Hox11/13b (Table S3; Figure S7C).

### Wnt signaling regulates Vasa expression

The coexpression of Wnt signaling ligands and a downstream target of Delta/Notch signaling, FoxY3, in Vasa-positive cells led us to test whether Wnt and Delta/Notch signaling may regulate germline factor expression in the sea star. To test whether ectopic activation of canonical Wnt signaling alters *P. miniata* germline factor gene expression, we treated embryos with the Wnt signaling activator LiCl over a developmental time course and measured effects on levels and distribution of the germline-specific transcripts Nanos and Vasa [35]. Embryos were cultured in fresh sea water and treated with 5mM LiCl for 24 hours, at three time points: one-day post-fertilization (Day 1), two-days post-fertilization (Day 2), or three-days post-fertilization (Day 3) (Figure 4A). At 24 hours after each treatment time point, Nanos and Vasa transcript levels were measured by whole-embryo quantitative RT-PCR (qPCR). Untreated embryos cultured and collected under the same conditions and timing were used as controls.

**Figure 4.**
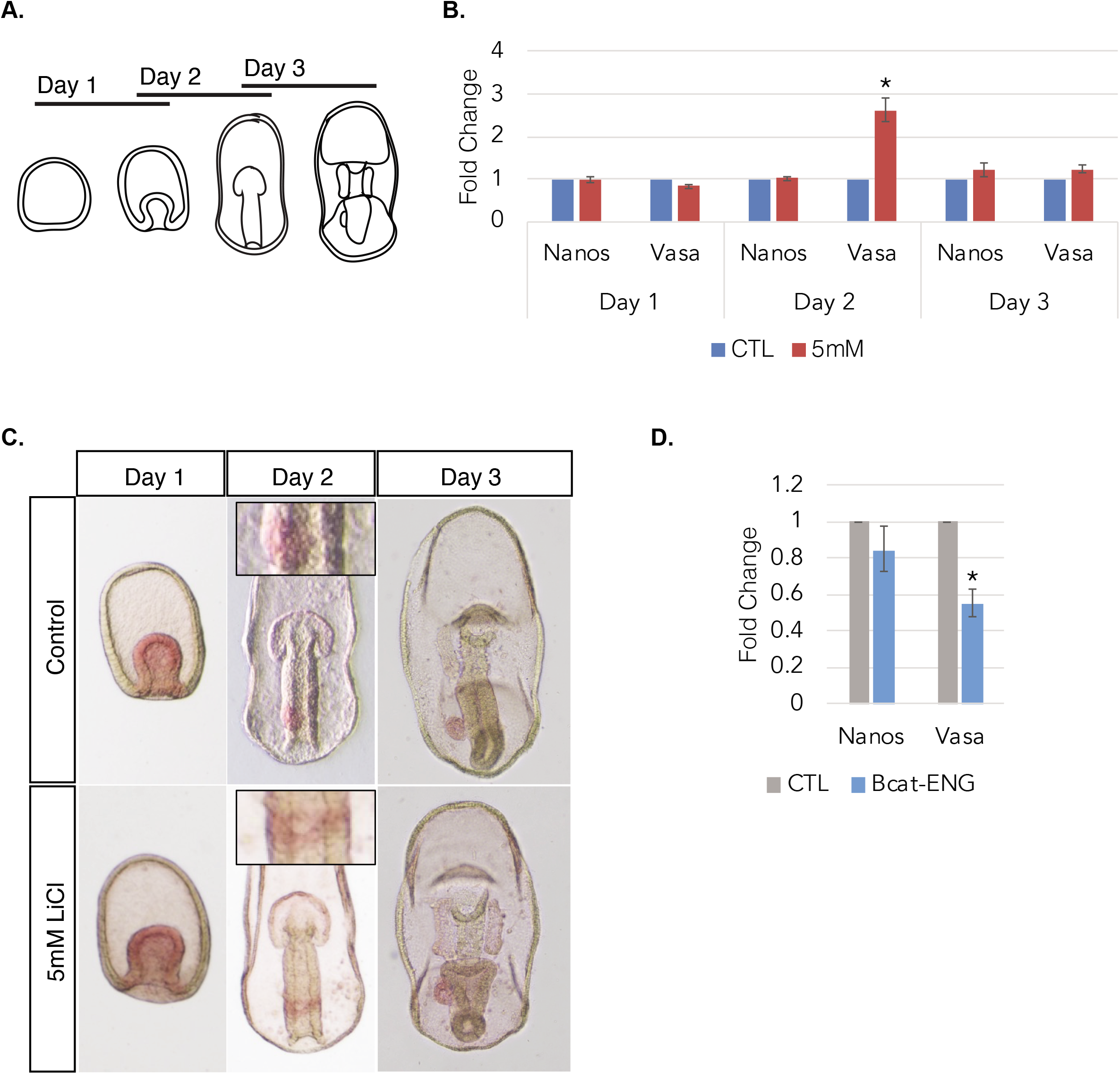
Wnt signaling regulates Vasa expression at gastrula stage. (A) Schematic showing experimental design. Embryos were treated with 5mM LiCl at one, two and three-days after fertilization for 24-hour incubation. (B) qPCR of Nanos and Vasa transcript abundance in control and LiCl treated embryos. Vasa transcript abundance is increased with treatment at day 2. Nanos expression is unaffected. (C) Vasa transcripts (red) accumulate on the left side of the gut in control embryos. Embryos treated with LiCl show Vasa expression as a band across the gut or loss of left-side restriction. (D) Injection with the transcriptional repressor construct ***β***-catenin/engrailed, results in a decrease of Vasa transcript abundance in gastrula stage embryos.

When Wnt signaling is upregulated, Vasa mRNA abundance is upregulated compared to controls; Nanos expression is unaffected (Figure 4B). Consistent with the qPCR results, Vasa mRNA expression expands with LiCl treatment at day 2 (Figure 4C). At this time point, the control embryos display Vasa staining as a rounded zone on the left edge of the gut. However, the LiCl-treated embryos display Vasa staining as a band from the left edge to the right edge of the gut. It is important to note that Vasa expression was not upregulated throughout the embryo but rather in a band along the gut. This result suggests that Wnt signaling regulates Vasa expression in the embryo, at gastrula stage.

To further test whether Vasa is a direct target of the canonical Wnt signaling pathway, we made use of a ***β***-catenin/Engrailed mRNA construct coding for a protein in which the transactivation domain on the ***β***-catenin C-terminus is replaced with the repression domain of the *Drosophila* Engrailed gene [34, 36]. Embryos injected with the transcriptional repressor construct showed a significant downregulation of Vasa transcripts at gastrula stage, while Nanos mRNA level was unaffected (Figure 4D). This result suggests that in gastrulae, Vasa may be a direct transcriptional target of canonical Wnt signaling. We have identified four putative HMG-box binding motifs recognized by ***β***-catenin and the transcription factor Tcf/Lef upstream of the Vasa promoter region that we will test in the future to determine whether Wnt regulation of Vasa is direct (5’-(T/A)ACAAAG-3’) [34, 37].

### Delta/Notch signaling regulates Nanos and Vasa expression

Delta/Notch signaling was found to regulate Nanos expression through the FoxY transcription factor, in somatic cells adjacent to the PGCs in the purple sea urchin [34]. In the sea star, the delta ligand is expressed in the same region of Nanos and Vasa expression, first in the invaginating gut of early gastrula, and then in the midgut region [27]. To test whether Delta/Notch may regulate germ cell factor gene expression in the sea star, we treated two-day old embryos with the Delta/Notch signaling inhibitor, DAPT [38]. Expression of both Nanos and Vasa is significantly decreased with DAPT treatment compared to controls. Expression of FoxY2 and FoxY3 is also decreased with DAPT treatment (Figure 5). *P. miniata* has three FoxY transcription factors, expression of all three is present at mid-gastrula, in the same region as Nanos and Vasa expression. By late gastrula stage, expression of FoxY1 and FoxY2 is lost (Figure S11). FoxY3 is expressed in the same domain as Nanos and Vasa at mid- and late gastrula stage (Figure S7A). Furthermore, we detect FoxY3 and Vasa coexpression at mid-gastrula by scRNA-seq (Figure S7B). To test whether FoxY3, downstream of Delta/Notch signaling, functions to regulate germline gene expression in the sea star, we used a morpholino approach to block translation of the protein. We do not see a significant change in mRNA transcript abundance for Nanos or Vasa with FoxY3 knockdown or knockdown of the other two FoxY transcription factors. We do note a significant upregulation in FoxY1, FoxY2, and FoxY3 transcripts with morpholino treatment suggesting that the three FoxY transcription factors behave as transcriptional repressors and regulate themselves through a negative feedback loop or that morpholino binding protects the transcripts from degradation (Figure S8). This data suggests that although FoxY3 is expressed in the same domain as Nanos and Vasa, it does not directly regulate their expression. The downstream Notch signaling effector, suppressor of hairless (SuH) is also downregulated with Delta/Notch inhibition (Figure 5). A morpholino or CRISPR/Cas9 approach targeting SuH would be a powerful test for whether Nanos and Vasa are direct transcriptional targets of Delta/Notch through SuH activity.

**Figure 5.**
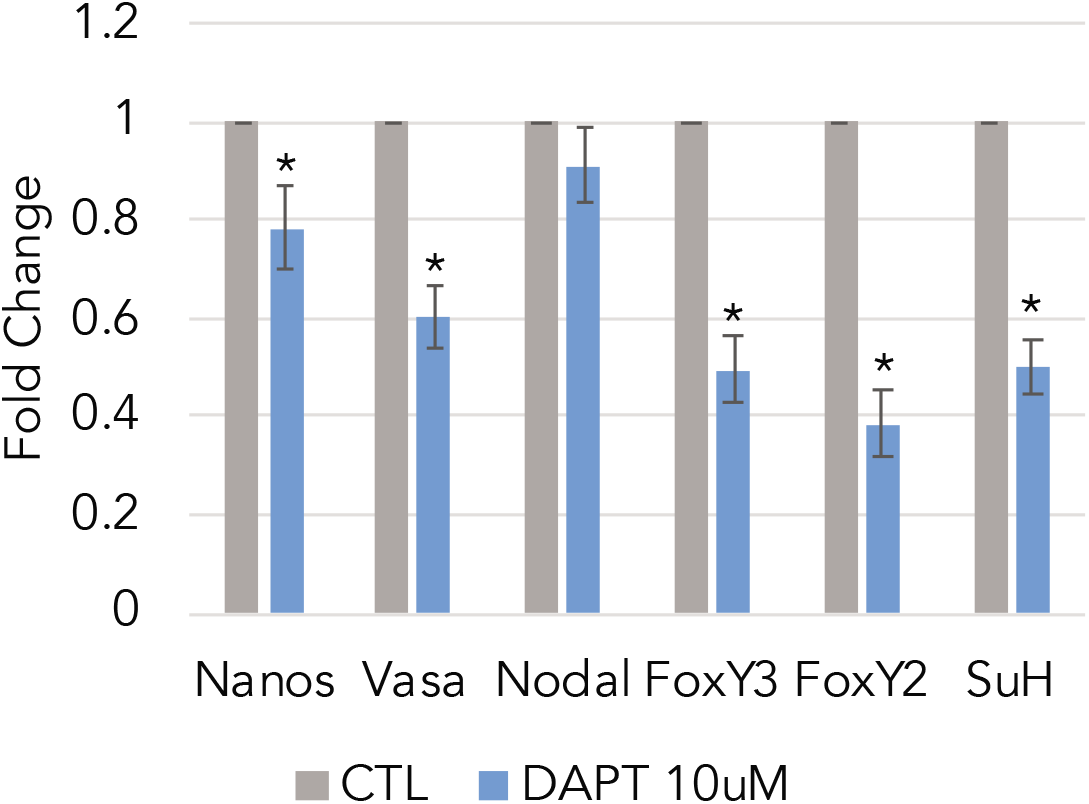
Delta/Notch regulation of germline genes. qPCR following incubation in 10uM DAPT at two-days post-fertilization. Nanos and Vasa expression decreases with DAPT treatment compared to controls.

### Analysis of cell interactions

The small molecule perturbation experiments and morpholino approaches help reveal the transcriptional mechanisms involved in germline factor expression in the sea star embryo. However, these experiments do not address the question of what or whether cell interactions are required for germline gene expression. We took advantage of the ability to dissociate embryos into single cells to test this premise. In having dissociated cells suspended in sea water we have inhibited signaling through cell contact and we have greatly diluted signaling ligands. Sea star embryos were cultured until hatched blastula stage at which point Nanos mRNA abundance is low [26]. Embryos were then dissociated and cultured for four hours, to early gastrula stage (Figure 6A). Dissociating cells at this stage in development allows us to ask the question of whether cell interactions are required to initiate Nanos expression and/or to maintain Vasa expression. Remarkably, Nanos transcript levels were significantly upregulated in dissociated cells compared to intact blastula and gastrula stage embryos whereas Vasa mRNA levels did not reach similar levels as in the intact gastrula. Nodal mRNA level was significantly decreased in dissociated cells compared to gastrula control, as might be expected when considering a positive feedback loop for Nodal in intact embryos [39, 40]. Dissociated cells showed upregulated Wnt8 expression and downregulated FoxY3 expression, consistent with a loss of cell contact and thus loss of Delta/Notch signaling. The level of Blimp1 mRNA, expressed in the same region as Nanos and Vasa, does not change with dissociation (Figure 6B). However, expression of endomesodermal genes known to correlate with nuclear ***β***-catenin activity in the sea star embryo are downregulated in dissociated cells (Figure S9C) [41]. Additionally, Nanos mRNA abundance continues to increase in dissociated cells cultured for up to 18 hours (Figure S9B).

**Figure 6.**
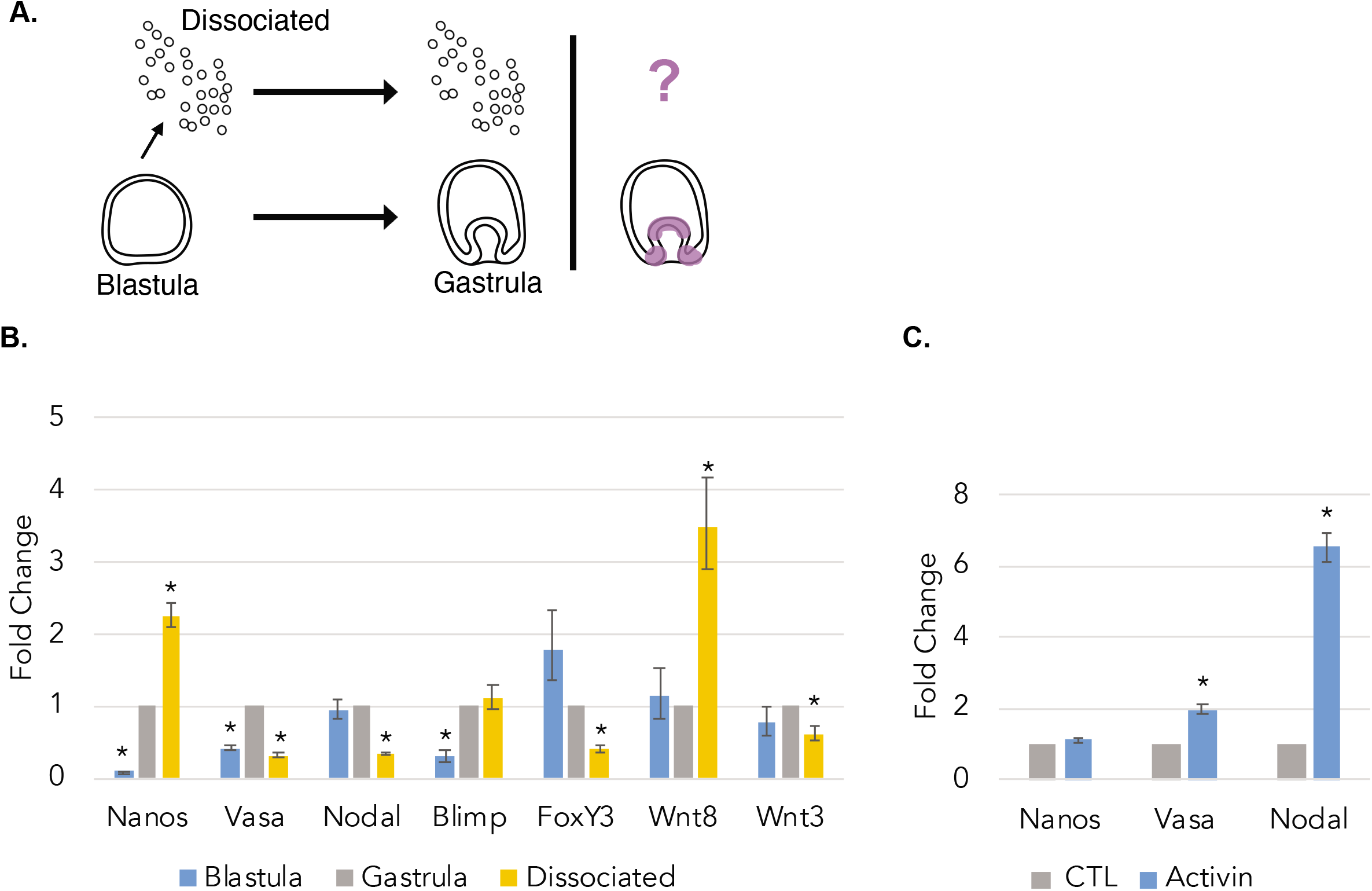
Analysis of cell interactions. (A) Schematic of experimental design. Embryos were cultured until hatched blastula stage at which point they were dissociated (D) and cultured for several hours until gastrula stage. (B) qPCR data showing change in mRNA transcript abundance with dissociation using intact gastrula stage as reference. Nanos expression is upregulated in dissociated cells compared to intact blastula and gastrula stage embryos. Vasa and Nodal expression in dissociated cells is downregulated compared to gastrula stage controls. (C)Treatment of dissociated cells with rhActivinAB results in Nodal upregulation. Nanos expression is unchanged compared to controls.

A potential explanation for the upregulation in Nanos transcripts is the drop in Nodal. Since Nodal represses Nanos mRNA accumulation, separation of cells from this interaction may minimize such a Nodal phenotype. To test the effect of Nodal signaling on dissociated cells, we treated dissociated cells with the Nodal signaling activator, recombinant human activin AB (rhActivinAB). In dissociated cells treated with rhActivinAB, Nodal expression is upregulated, the spike in Nanos mRNA seen with dissociation is lost, and Vasa expression is now upregulated (Figure 6C). We previously reported an inhibitory effect of Nodal on Nanos expression [7]. A loss of Nanos expression with rhActivinAB treatment suggests that Nodal acts directly on Nanos expressing cells to regulate Nanos expression. At this time, it is unclear whether the upregulation of Nanos transcripts in dissociated cells compared to controls is due to a greater number of cells expressing Nanos or a small population of cells greatly upregulating Nanos expression. The former would indicate that Nanos expression is cell autonomous and Nodal plays an essential role in restricting Nanos expression throughout the embryo.

## Discussion

### Inductive signals in sea star germ cell specification

Our current understanding of the inductive mechanism for germ cell specification comes from studies performed using mouse, cricket, and axolotl embryos. Identifying activating and inhibitory cell signaling pathways and their targets is essential to understand germline induction across different species, how these processes may be disrupted to effect fertility, and how they can be utilized for gamete development studies in vitro [42, 43]. The sea star possesses many strengths as a model organism for study of the inductive mechanism including access to greater numbers of oocytes, and manipulation and testing of early embryos at different points in the specification process. Here we present a role for Wnt and Delta/Notch signaling in inductive germline specification of sea star embryos through regulation of the germline genes, Nanos and Vasa. We propose that both Wnt and Delta/Notch signaling regulate Nanos and Vasa expression at gastrula stage. These pathways could be active concurrently with Nodal restriction which originates from a gradient on the right side of the embryo. While Delta/Notch signaling affects both Nanos and Vasa expression, Vasa alone seems to be under Wnt/***β***-catenin regulation. This suggests that although Nanos and Vasa are expressed in the same regions, and similarly repressed by Nodal, how that expression pattern comes to be is distinct. Further investigation is necessary to uncover whether both genes are direct transcriptional targets of Delta/Notch and Wnt. We also report a difference in Nanos and Vasa regulation through dissociation experiments. In this setting, with cell signaling removed, Nanos transcript abundance increases and continues to do so with time, while Vasa transcript abundance does not. This experiment reveals that activating signals are essential for initiating and maintaining Vasa expression in the embryo. Nanos does not seem to face the same regulation. Future studies will reveal whether Nanos expression is cell autonomous and if cells throughout the early embryo are capable of expressing Nanos in the absence of a repressor. It is possible that particularly for Nanos, Nodal repression is crucial in promoting proper development of the germline and the embryo as a whole.

At the sea star mid-gastrula stage, Vasa transcript is expressed more broadly than in the future germline, this contrasts its expression in the sea urchin where it is restricted to the small micromeres in gastrulae and the mouse where it is expressed only after the PGCs have colonized the embryonic gonad [44, 45]. This is a distinct strategy of cellspecific germ cell factor accumulation in which expression is spread broadly then cleared from cells to yield highly localized expression. In the sea star, Vasa protein is also under strict control through selective post-translational degradation by Gustavus, an E3 ligase [46]. In the future, investigating how Nanos protein localization is regulated will inform us whether these transcripts are also under distinct regulation at the protein level.

During gastrulation, Vasa and Nanos mRNA expression is present in the archenteron. And in larvae, Nanos and Vasa expression is present in the PE as well as the anterior pouches [7]. We detected Nanos and Vasa transcripts as well as FoxY3 at the tip of the archenteron (mid-gastrula cluster 3) (Figures S6G-I and S7). What role these transcripts, particularly Nanos and Vasa, have in this region is unknown. A notable difference in germline development in the sea urchin compared to sea star is that a PE structure is absent. Instead, the germline determinants, including Nanos and Vasa, accumulate in the small micromeres and become incorporated into both coelomic pouches before becoming restricted to the left pouch [44]. Future studies are essential to determine whether these cells expressing Nanos and Vasa at the tip of the archenteron have a function in germline development conserved in the sea urchin embryo.

### Distinct mechanisms of germ line specification in two echinoderm species

In comparable sea urchin developmental stages, more cell states and regions of the embryo are partitioned in distinct clusters than in the sea star. For example, at early blastula stage in the sea urchin, skeleton, pigment, neural cell states, and distinct oral ectoderm cell states are present. By early gastrula, scRNA-seq identified six ectodermal cell states and three neural cell states [12]. Of note, the skeletogenic and pigment cell lineages are mesodermal cell types of the sea urchin, absent in the sea star embryo. The sea star (*P. miniata*) and sea urchin (*P. lividus*) have been compared to assess heterochronies in marker gene expression, revealing a later onset of maternal to zygotic transition in the sea star compared to this sea urchin [47]. It is possible that this temporal difference plays a role in the diversity of cell states present at the early developmental stages when comparing the two species. An alternative explanation is that cell fates in the sea star rely more on cell signaling for fate decisions than in the sea urchin, and the sea star embryonic cells are less committed than in the comparable stage of a sea urchin.

Echinoderms offer an opportunity to investigate two modes of germ cell development in closely related species and uncover evolutionary differences. Nanos and Vasa eventually localize to putative PGC precursors in both species (the small micromeres in sea urchins and in the PE in the sea star), yet their embryonic expression patterns are vastly different. Early in sea urchin embryogenesis, Nanos and Vasa expression is restricted to the small micromeres [44]. In the sea urchin, Nanos transcription is regulated by direct action of ***β***-catenin in the small micromeres or the Delta/Notch transcription factor, FoxY, in the adjacent somatic Veg2 mesoderm cells [34]. In the sea star, Vasa mRNA expression is seen earlier than Nanos, present in the oocyte, in the vegetal pole of blastulae, and later with Nanos in the hindgut and archenteron regions [7, 8, 26]. Similar to the sea urchin, Delta/Notch regulation of Nanos is also present in the sea star (Figure 5). However, we propose that the FoxY transcription factors present in the sea star are not transcriptional regulators of Nanos. This could point to an evolutionary shift between the two species where FoxY plays a role in germ line specification but its transcriptional targets have changed. A difference in Wnt signaling effect is also evident. Nanos expression in the small micromeres is regulated by ***β***-catenin in the sea urchin. Although we do not report an effect on Nanos expression through Wnt signaling activation or repression in the sea star, analysis of different stages of development as well as the Nanos regulatory region are necessary to reach this conclusion. Notably, removal of the micromeres, the parent cells of the small micromeres, results in adults with developed gametes [48]. Whether Nanos expression, downstream of FoxY, in the Veg2 cells plays a role in reconstituting the germline remains an interesting question and could point to an auxiliary, inductive mechanism present in the sea urchin. Testing effects of cell signaling perturbation in micromere-deleted embryos will help decipher this process.

### Evolution of germ line specification

Evidence argues the inductive mode (epigenesis) in metazoans such as sponges, jellyfish, and hydra is the ancestral mechanism of germ cell specification [1]. Discovery of an inductive mode for germline specification in the sea star, an echinoderm species, presents an opportunity for comparison with the mouse, a mammalian species for which we have the most comprehensive description of the inductive mechanism (Figure 7). Mouse PGCs acquire their fate in response to signaling. They arise from mesodermally differentiating cells. Single-cell analysis revealed that early germ cells express homeobox genes, including Hoxb1 and Hoxa1, and the mesodermal marker brachyury, at the same level as their somatic cell neighbors. The early germ cells then initiate transcriptional repression of homeobox genes which remain highly expressed in the neighboring somatic cells as the embryo develops [49]. Here we see a similarity in the sea star where we see coexpression of Hox11/13b and brachyury with Nanos-positive cells (Figure S7D). In the mouse, expression of germline determinants with mesodermal markers is resolved by repression of the mesodermal markers in the PGC precursor cells; Stella-positive cells repress homeobox genes [49]. In the sea star, apart from restricting Nanos and Vasa transcript expression, repression of mesodermal markers in Nanos- and Vasa-positive cells remains an important step in germline specification which has yet to be identified.

**Figure 7.**
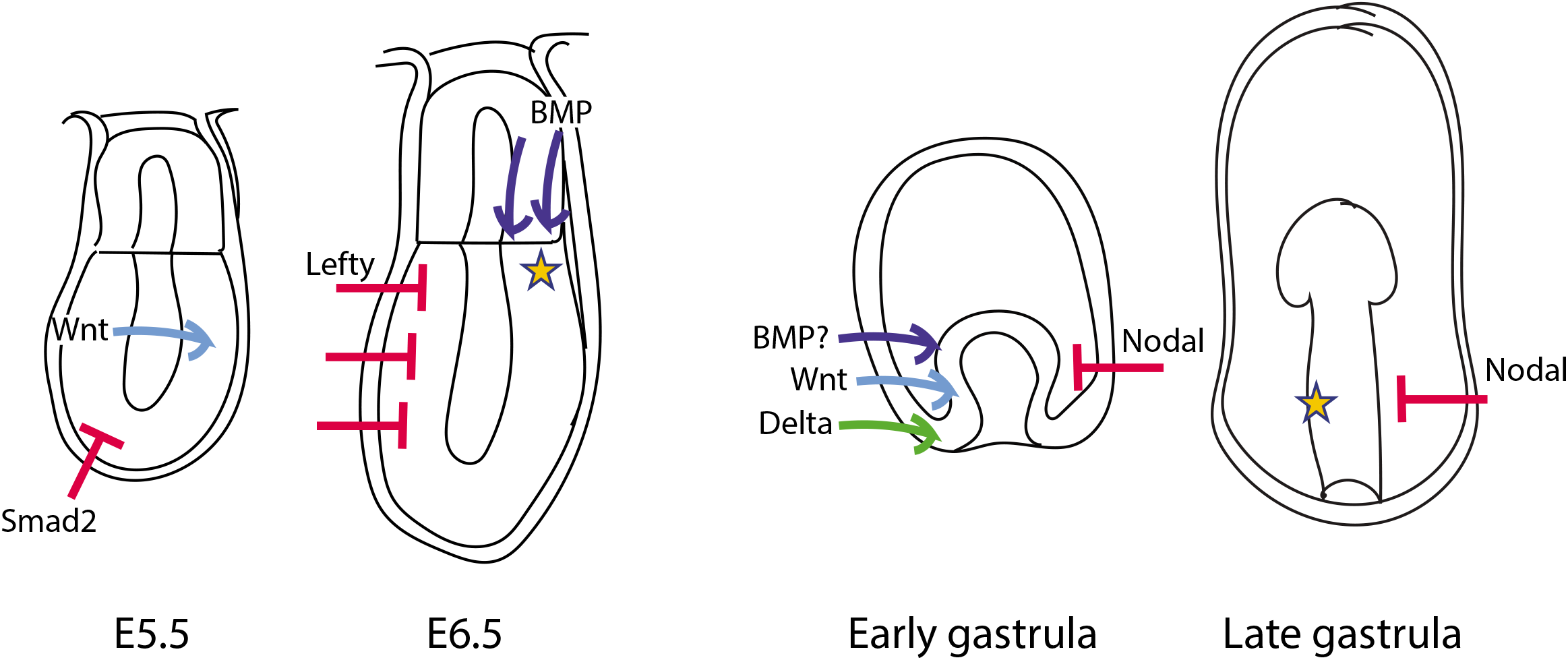
Inductive mechanism in mouse and sea star embryos. Model highlighting similarities and differences between inductive germline development in mouse and sea star. Early inductive Wnt signaling conserved between mouse and sea star, with a role for Delta/Notch identified in the sea star. Nodal signaling has a conserved role in restricting germline gene expression in both species. Preliminary data suggests BMP may have a conserved role in activating germline gene expression.

In the mouse embryo, Wnt3 and Nodal signaling serve as early regulators of PGC specification. These early signals give the epiblast cells competence to respond to BMP signaling, which regulates expression of germline determinants like Blimp1 [3, 6]. This report of Wnt regulation of Vasa expression suggests that Wnt signaling may have an ancestral, conserved role in inductive germline formation. Nodal also has a role in restricting the germline in both mouse and sea star. Further studies are necessary to identify which Wnt ligand is responsible for regulating germ cell development in the sea star. The early vegetal expression of Wnt3, Wnt16, Wnt8, and Frizzled1 receptor makes them potential candidates for testing their role in germ cell specification [25]. Following the inductive mechanism, PGC specification is a sequential process. This is evident in the sea star through sequential restriction of germline gene expression first to the dorsal side and later to the left side [7]. In the mouse, Wnt and Nodal signaling effects are required before a response to BMP signals is seen [3]. An important future direction is the examination of BMP signaling in the sea star. BMP signaling has a conserved role in germline development by regulating expression of germline genes in mouse, cricket, axolotl (*Ambystoma mexicanum*) and the acorn worm (*Ptychodera flava*) [3, 4, 50, 51]. Preliminary data suggests that BMP may have a role in activating Vasa mRNA expression in the early gastrula (Figure S10). Further investigation of sea star embryos will uncover the sequence of signaling events required for proper development of the germ cells and whether these signaling events have been conserved through evolution.

## Supporting information

Foster et al

## Supplementary Figure Legends

**Supplementary Figure S1. Identification of cell states across early sea star development.** UMAP visualization of six integrated datasets, separated by embryonic stage: 8-hpf (868 cells), 10-hpf (1,318 cells), 14-hpf (2,448 cells), blastula (7,272 cells), early gastrula (3,349 cells), and mid-gastrula (10,448 cells).

**Supplementary Figure S2. Marker gene expression at blastula stage.** Feature plots showing expression of ectodermal, mesodermal and endodermal marker genes at blastula stage. Average gene expression level displayed by color intensity.

**Supplementary Figure S3. Marker gene expression at mid-gastrula stage.** Feature plots showing expression of ectodermal, mesodermal and endodermal marker genes at mid-gastrula stage. Average gene expression level displayed by color intensity.

**Supplementary Figure S4. Wnt signaling component expression.** (A) Dotplot of Wnt signaling pathway component expression at blastula stage. Coexpression of Wnt3, Wnt16, and Frizzled receptor (Frizz) seen in cluster 2. (B) Dotplot of Wnt signaling pathway component expression at mid-gastrula stage.

**Supplementary Figure S5. MifL1-enriched cluster expression.** Dotplot showing expression of sea urchin PMC marker genes Mt1-4 and Fos in cluster 10, MifL1-enriched cluster, at mid-gastrula stage.

**Supplementary Figure S6. Vasa, Nanos, and FoxY3 expression.** Violin plots showing Nanos, Vasa, and FoxY3 expression per cluster in blastula stage (A-C) and mid-gastrula stage (G-I). Feature plots showing expression of Nanos, Vasa, and FoxY3 in blastula stage (D-F) and in cluster 8 of mid-gastrula stage (J-L).

**Supplementary Figure S7. Vasa and FoxY3 expression.** (A) Expression of FoxY3 in the developing embryo shown by in situ hybridization. (B) Vasa expression in 395 cells of mid-gastrula cluster 8 (hindgut) shown in red (top left). FoxY3 expression in 221 cells of cluster 8 shown in green (top right). Merge showing co-expression of Vasa and FoxY3 in 167 cells (bottom left). Color scale for expression level (bottom right). (C) Heatmap comparing expression in cells co-expressing Vasa and FoxY3 to those that do not co-express Vasa and FoxY3 in cells of cluster 8 of mid-gastrula stage (hindgut). (E) Violin plot of Hox11/13b and Brachyury expression in Nanos negative and Nanos positive cells of mid-gastrula cluster 8.

**Supplementary Figure S8. FoxY morpholino knockdowns.** qPCR data showing morpholino knockdown of FoxY1, FoxY2, or FoxY3 does not affect Nanos and Vasa mRNA abundance.

**Supplementary Figure S9. Changes in transcript abundance with dissociation.** (A) qPCR data showing change in mRNA transcript abundance with dissociation using intact blastula embryos as control. (Right panel is zoomed in.) (B) Nanos transcript abundance increases with time in dissociated cells compartment to intact gastrula. Embryos and dissociated cells were cultured for 3 hours (T1) and 18 hours after blastula hatching (T2). (C) Transcript abundance of gut specific, Wnt target genes is decreased in dissociated cells compared to intact gastrula.

**Supplementary Figure S10. BMP4 treatment in early sea star embryos.** In situ hybridization for Vasa mRNA expression in embryos treated with 50ng/mL or 100ng/mL recombinant human BMP4 at one, two and three days after fertilization for 24-hour incubation. Controls were incubated with 0.1% HCl BSA.

**Supplementary Figure S11. Nanos and FoxY gene expression.** Whole mount in situ hybridizations for Nanos and FoxY transcription factors during early sea star embryogenesis. Nanos3 expression is seen in mid-gastrula at the tip of the archenteron and the hindgut region. Nanos3 expression persists as a ring in the gut in late gastrula and becomes restricted to the PE; some expression is detected in the anterior pouches. Nanos2 is expressed transiently as a ring around the blastopore in mid-gastrula. FoxY1 and FoxY2 are also transiently expressed at mid-gastrula stage. FoxY3 is expressed in the same domain as Nanos3 at mid- and late gastrula stages.

## Material and Methods

### Embryo culture and dissociation

Adult *Patiria miniata* animals were collected by either Peter Halmay or Josh Ross off the Californian coast. Embryos were cultured essentially as described previously (Fresques et al., 2016). Embryos were cultured in filtered (0.2micron) sea water collected at the Marine Biological laboratories in Woods Hole MA, until the appropriate stage for dissociation. All embryos used in the study resulted from mating of one male and one female. Multiple fertilizations were initiated in this study and timed such that the appropriate stages of embryonic development were reached at a common endpoint. The embryos were then collected and washed twice with calcium-free sea water, and then suspended hyalin-extraction media (HEM) for 10-15 minutes, depending on the stage of dissociation. When cells were beginning to dissociate, the embryos were collected and washed in 0.5M NaCl, gently sheared with a pipette, run through a 40micron Nitex mesh, counted on a hemocytometer, and diluted to reach the appropriate concentration for the scRNA-seq protocol. Equal numbers of embryos were used in each time point and at no time were cells or embryos pelleted in a centrifuge (Oulhen et al., 2019).

### In situ hybridization

DIG-labeled RNA probes were made with a Roche DIG probe synthesis kit as described previously (Fresques et al., 2016). Probe-hybridized embryos were developed with NBT+BCIP for purple. Embryos were incubated with probe for 1 week and were developed essentially as described previously (Fresques et al., 2016).

### Pharmacological perturbation

Embryos were incubated with 5 mM LiCl, 10uM DAPT, 106ng/uL recombinant human activin AB, or recombinant human BMP4 at 50 or 100ng/mL, at the developmental time points indicated.

### Morpholino injection

Oocytes were injected with 1.0 mM of morpholino (at stock concentration, Gene-Tools) in injection solution: 10% glycerol and 1 mM Texas Red Dextran or 0.83 mM FITC Dextran (Molecular Probes). After incubation for 1 day at 16°C, injected fluorescent oocytes were selected and matured with 1-methyl adenine (3.0 μM Acros Organics). Eggs were fertilized with diluted sperm, washed and incubated in filtered sea water at 16°C until they developed to the desired embryonic stage.

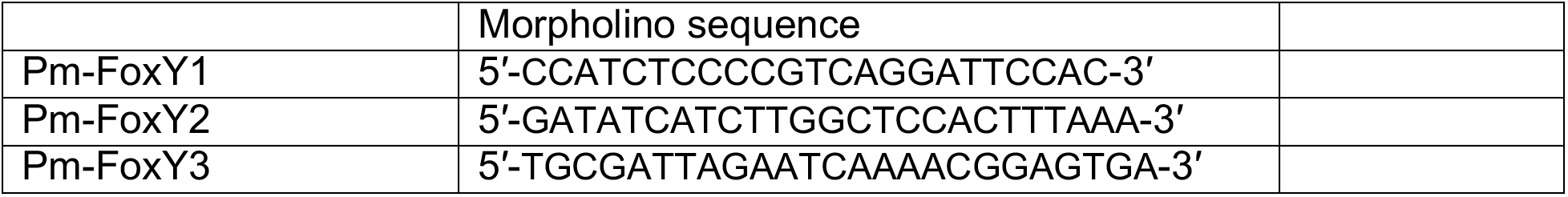

### Repressor construct injection

Oocytes were injected with 1.0mM of the ***β***-catenin/engrailed mRNA construct, cultured and collected at two-days post-fertilization for qPCR analysis.

### qPCR analysis

RNA was extracted from ~50 embryos in each desired stage by using the RNeasy Micro kit (Qiagen). A reverse transcription reaction was performed using MMLV Reverse Transcriptase (Promega) or M-MuLV Reverse Transcriptase (Thermoscientific, Maxima first-strand cDNA synthesis kit). Sybr green was used for qPCR analysis (ThermoFisher). Experiments were carried out either in duplicate or in triplicate, and ubiquitin or 18 s was used to normalize RNA levels between samples. All data are represented as fold-change relative to the standards. A one-tailed t-test was used to calculate significance. Error bars indicate ±1s.d.

### Single cell RNA sequencing

Single cell RNA sequencing: Single cell encapsulation was performed using the Chromium Single Cell Chip B kit on the 10x Genomics Chromium Controller. Single cell cDNA and libraries were prepared using the Chromium Single Cell 3’ Reagent kit v3 Chemistry. Libraries were sequenced by Genewiz on the Illumina Hiseq (2×150 bp paired-end runs). Single cell unique molecular identifier (UMI) counting (counting of unique barcodes given to individual transcript molecules), was performed using Cell Ranger Single Cell Software Suite 3.0.2 from 10X Genomics. The custom transcriptome reference was generated from *P. miniata* assembly V2.0 (echinobase.org) using CellRanger mkref. The annotation for PmNanos was manually edited to account for a longer 3’UTR. Duplicate blastula and gastrula stage libraries were aggregated using the cellranger aggr function. Cellranger gene expression matrices were further analyzed using the R package Seurat v 3.1.4 (Butler et al., 2019). Cells of 8hpf to 14hpf stages with at least 1000 and at most 6000 genes (features), and cells with at least 1000 and at most 2500 genes in blastula to mid-gastrula stages were included in downstream analysis. Individual datasets were normalized by scaling gene expression in each cell by total gene expression and then log transformed. The top 2000 highly variable genes across the datasets were then used to integrate the datasets. Individual time point datasets were integrated using the Seurat toolkit Harmony to identify conserved cell populations across the datasets. The clustering parameters used were dimensions: 20, resolution: 1.0. Cluster markers were found using FindConservedMarkers and FindMarkers functions.

