## Supplementary material for "Distinct mechanisms of germ cell factor regulation for an inductive germ cell fate": Foster et al

**Figure S1. Identification of cell states across early sea star development**

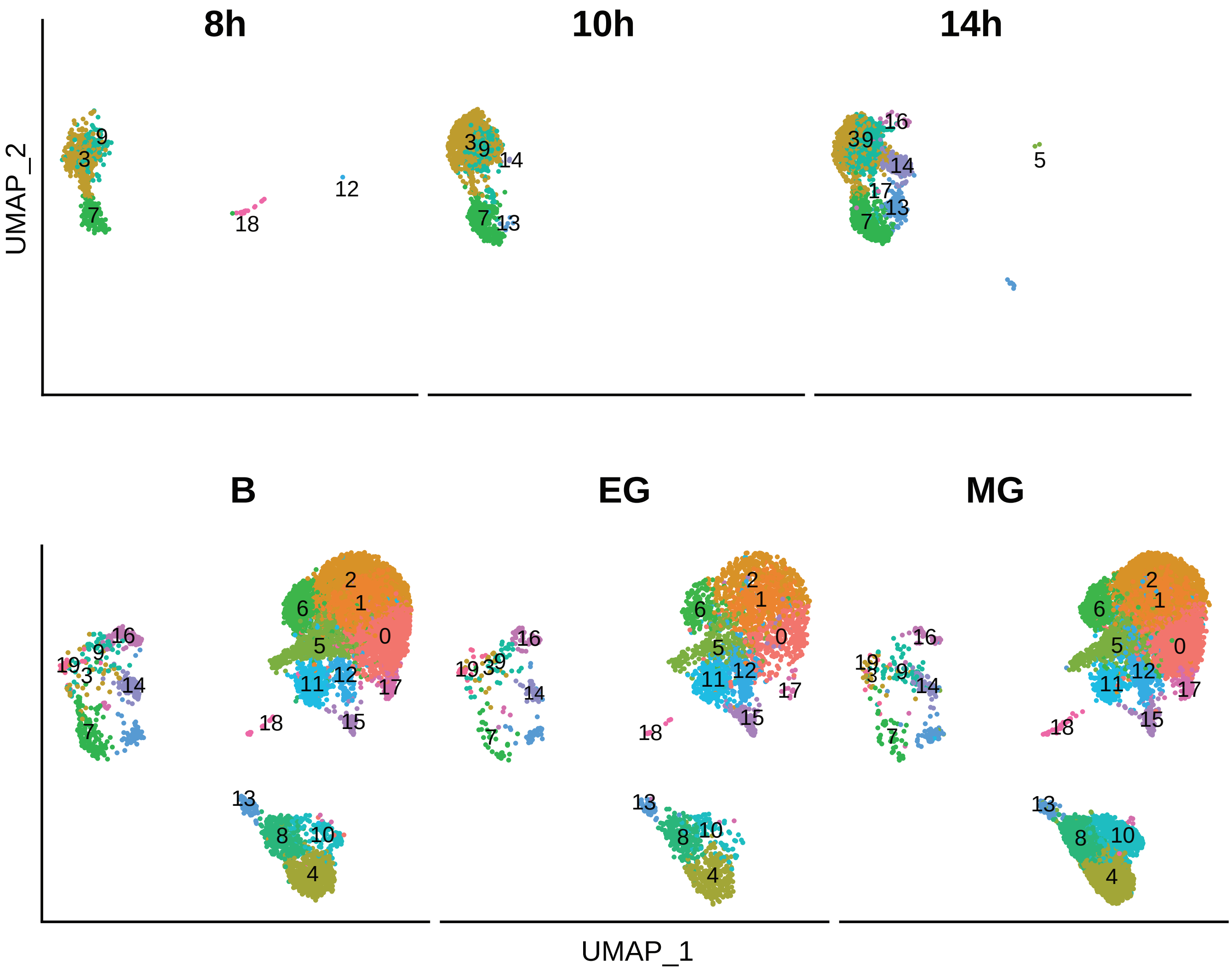

**Figure S2. Marker gene expression at blastula stage**

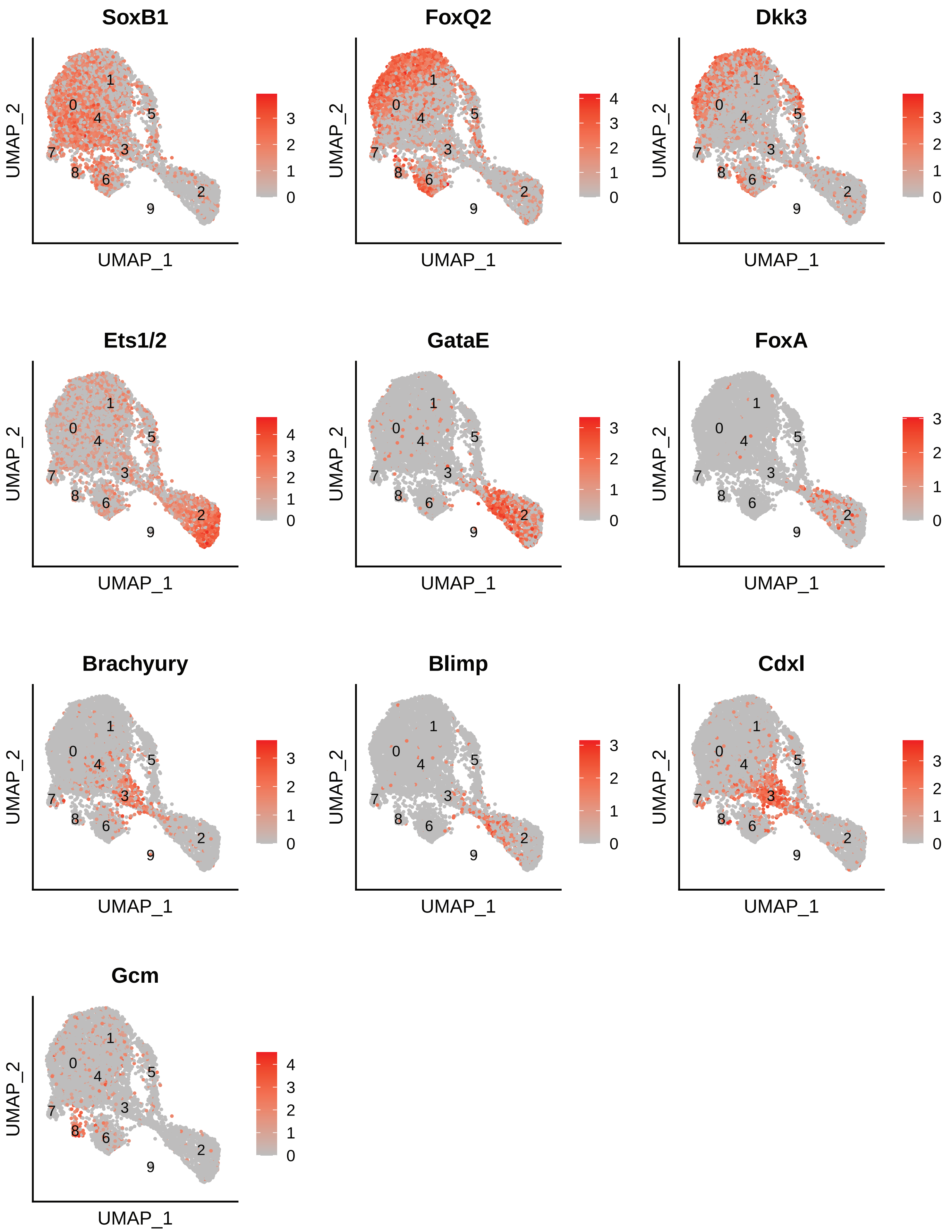

**Figure S3. Marker gene expression at mid-gastrula stage**

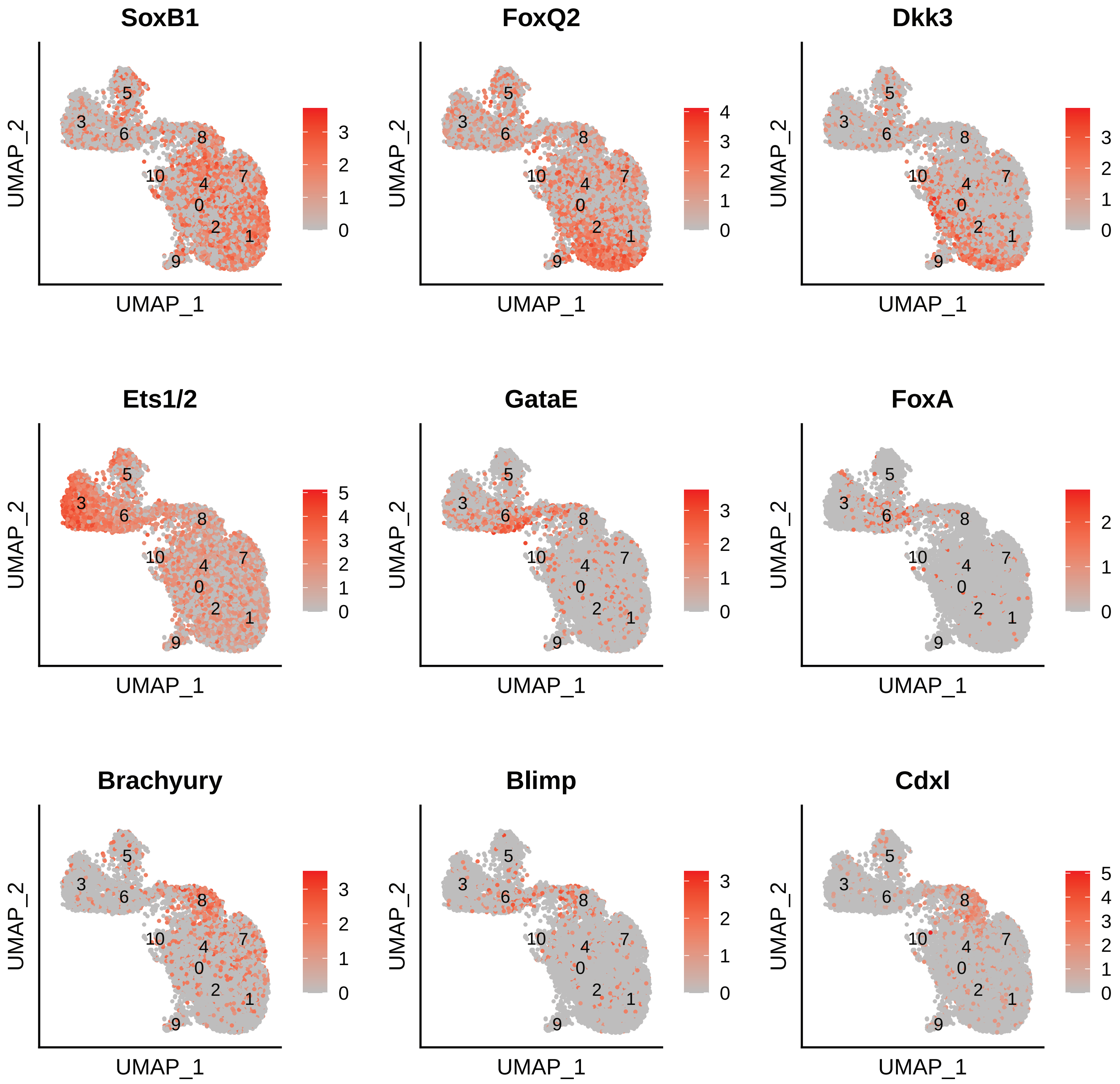

### Figure S4. Wnt signaling component expression

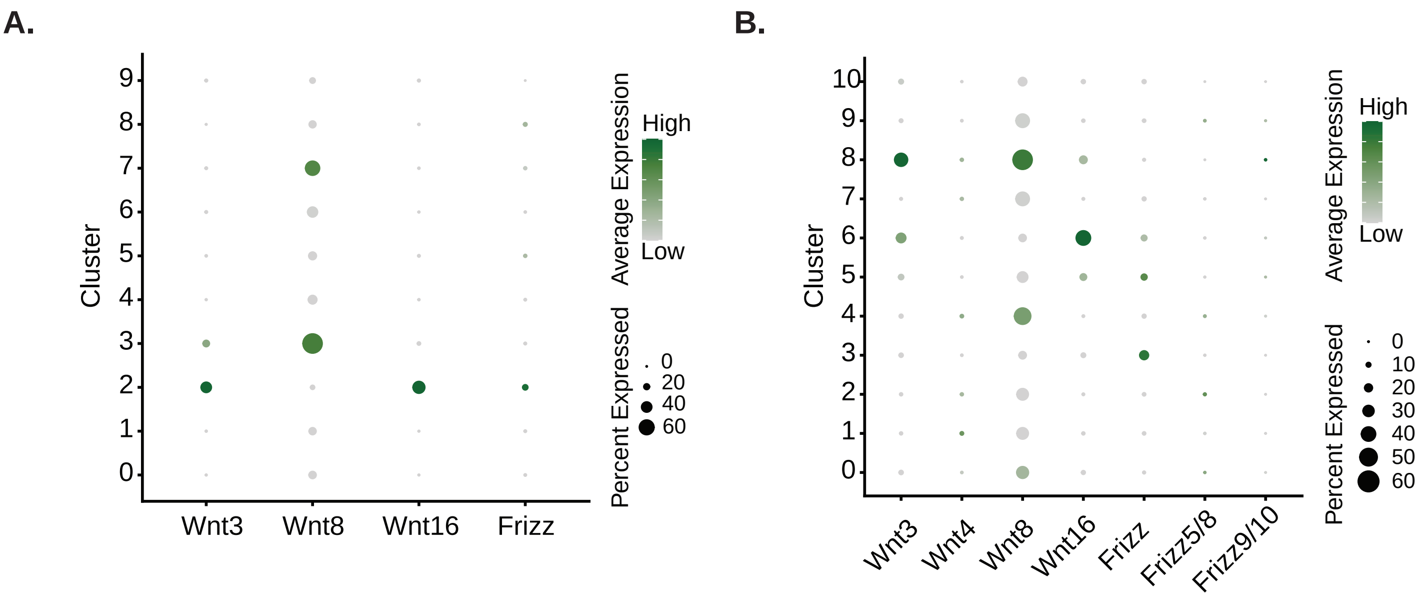

**Figure S5. MifL1-enriched cluster expression**

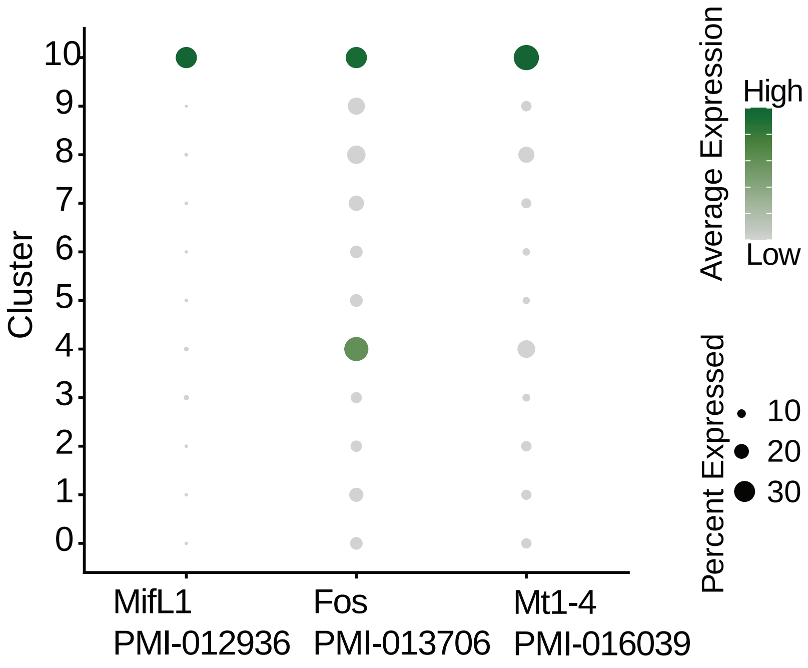

**Figure S6. Germline gene expression**

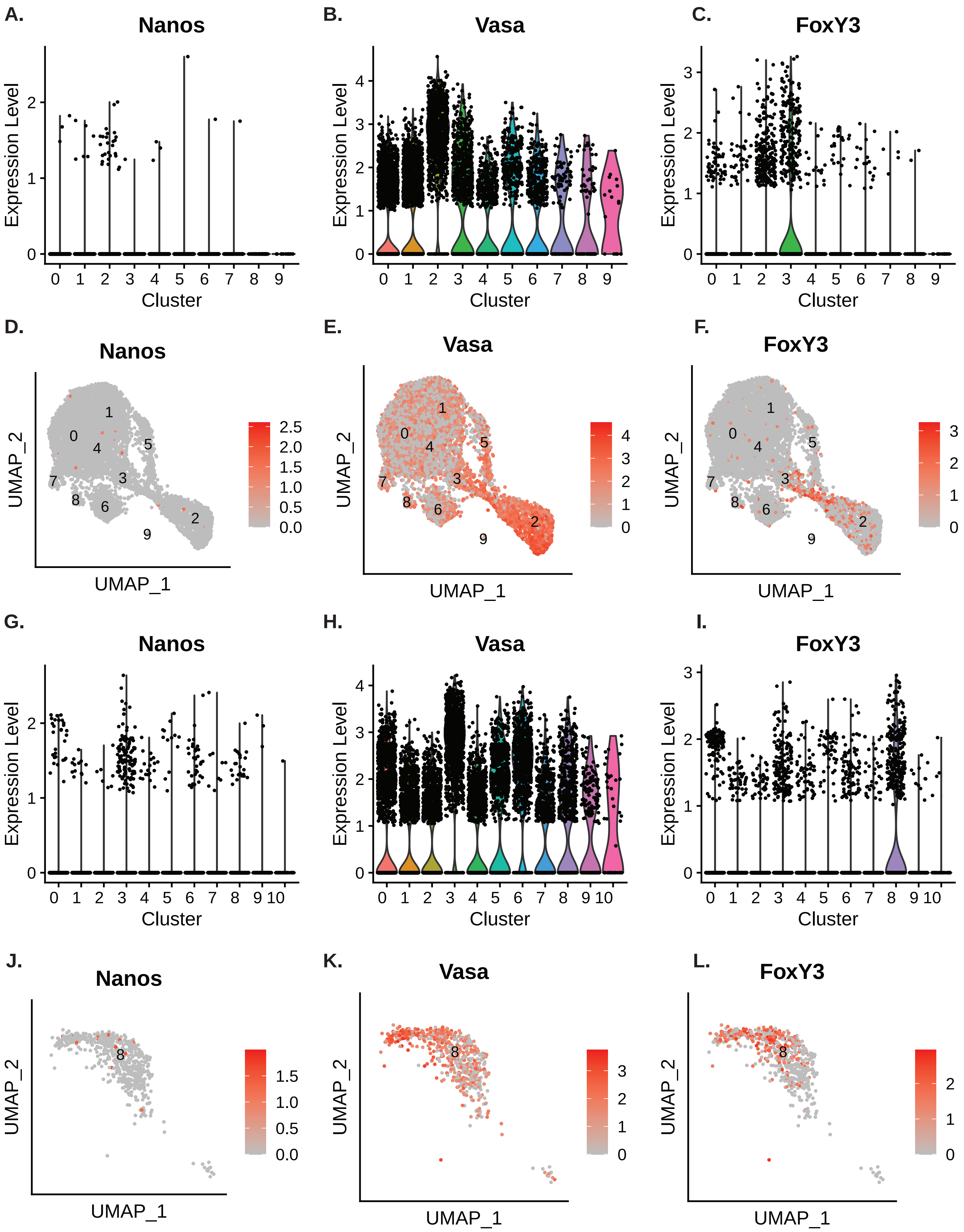

Figure S7. Vasa and FoxY3 expression

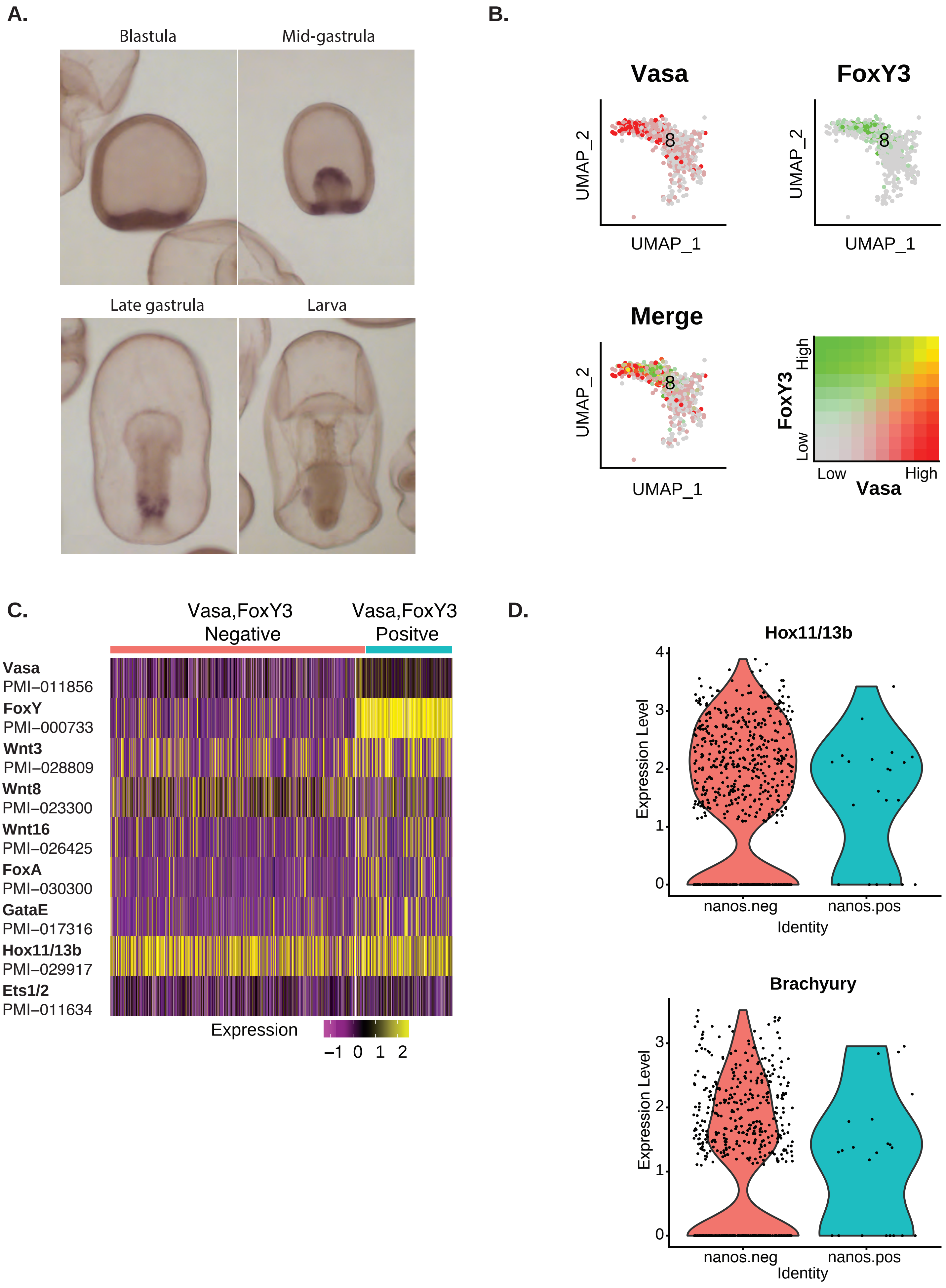

**Figure S8. FoxY transcription factor knockdowns**

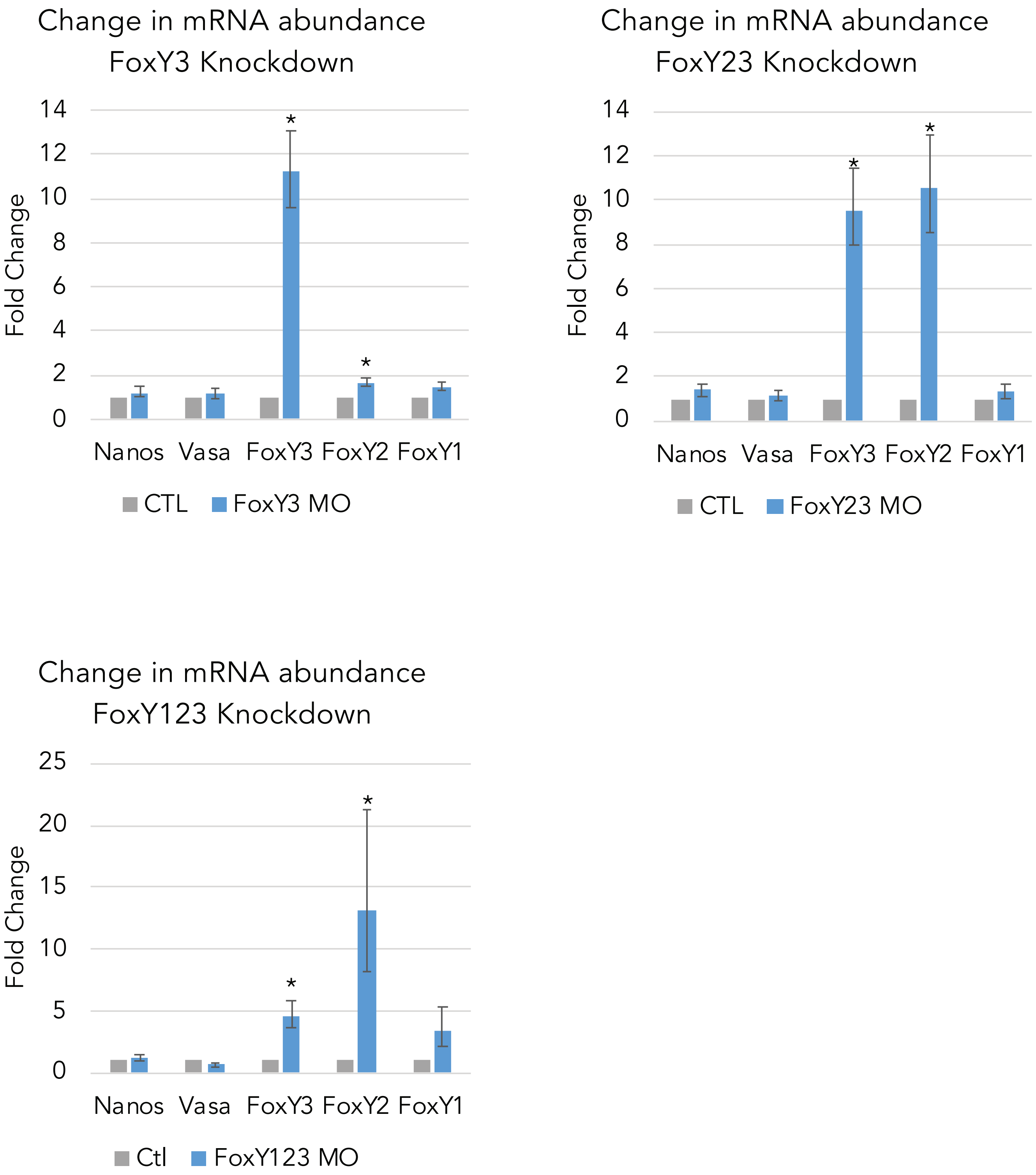

**Figure S9. Changes in transcript abundance with dissociation**

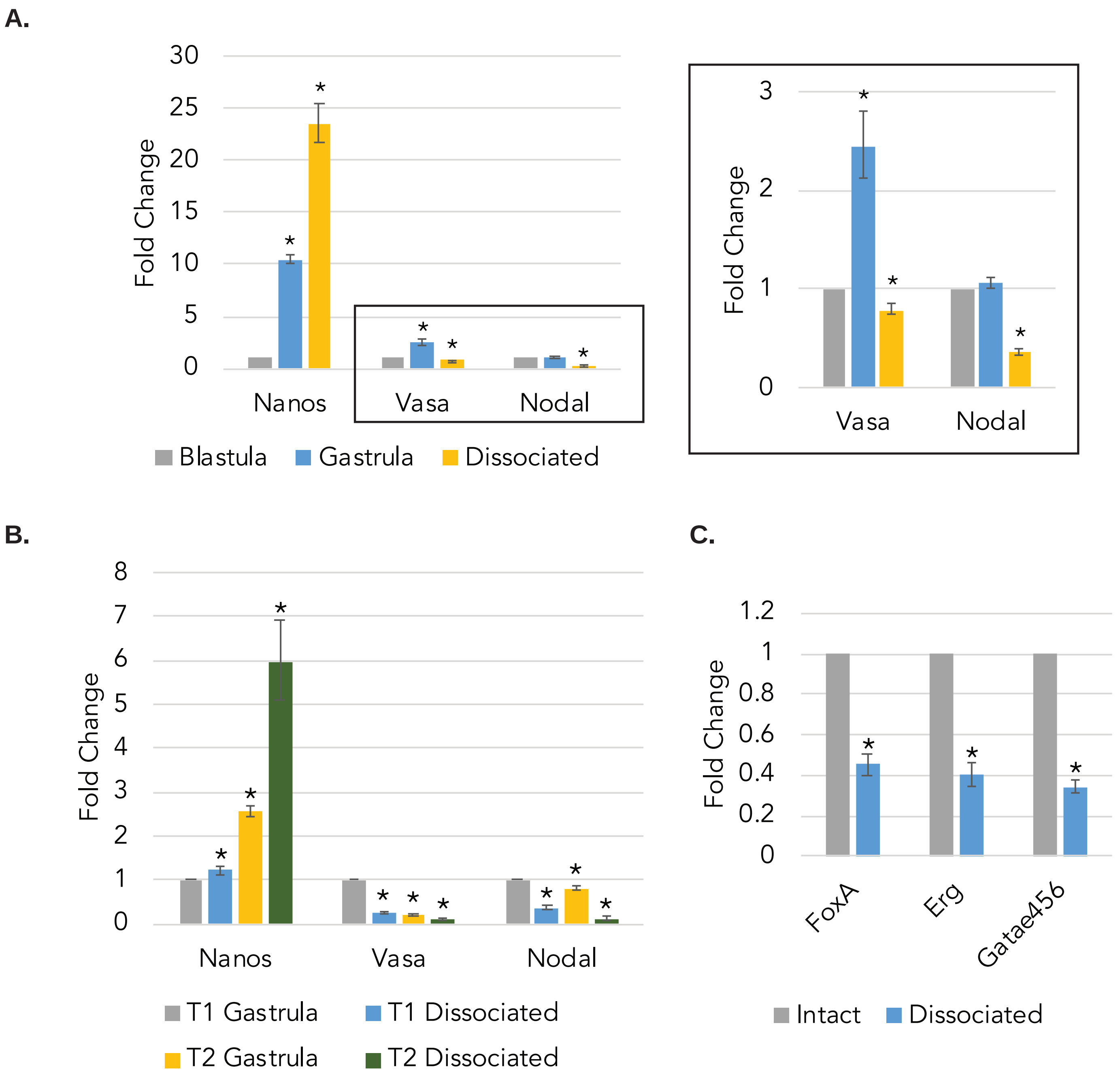

**Figure S10. BMP4 treatment of early embryos**

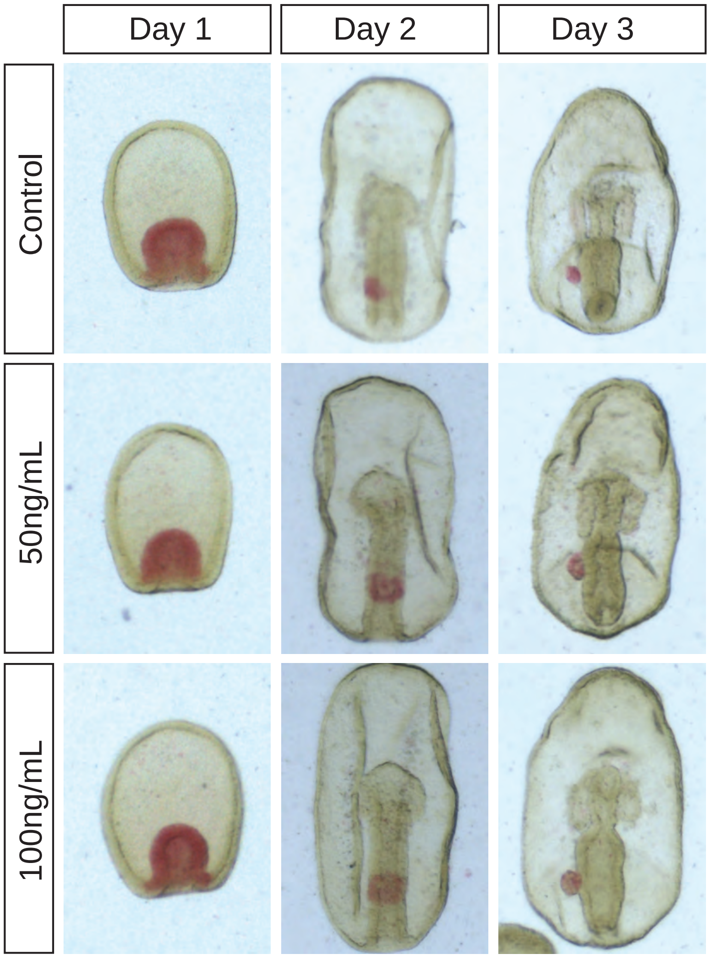

**Figure S11. Nanos and FoxY gene expression**

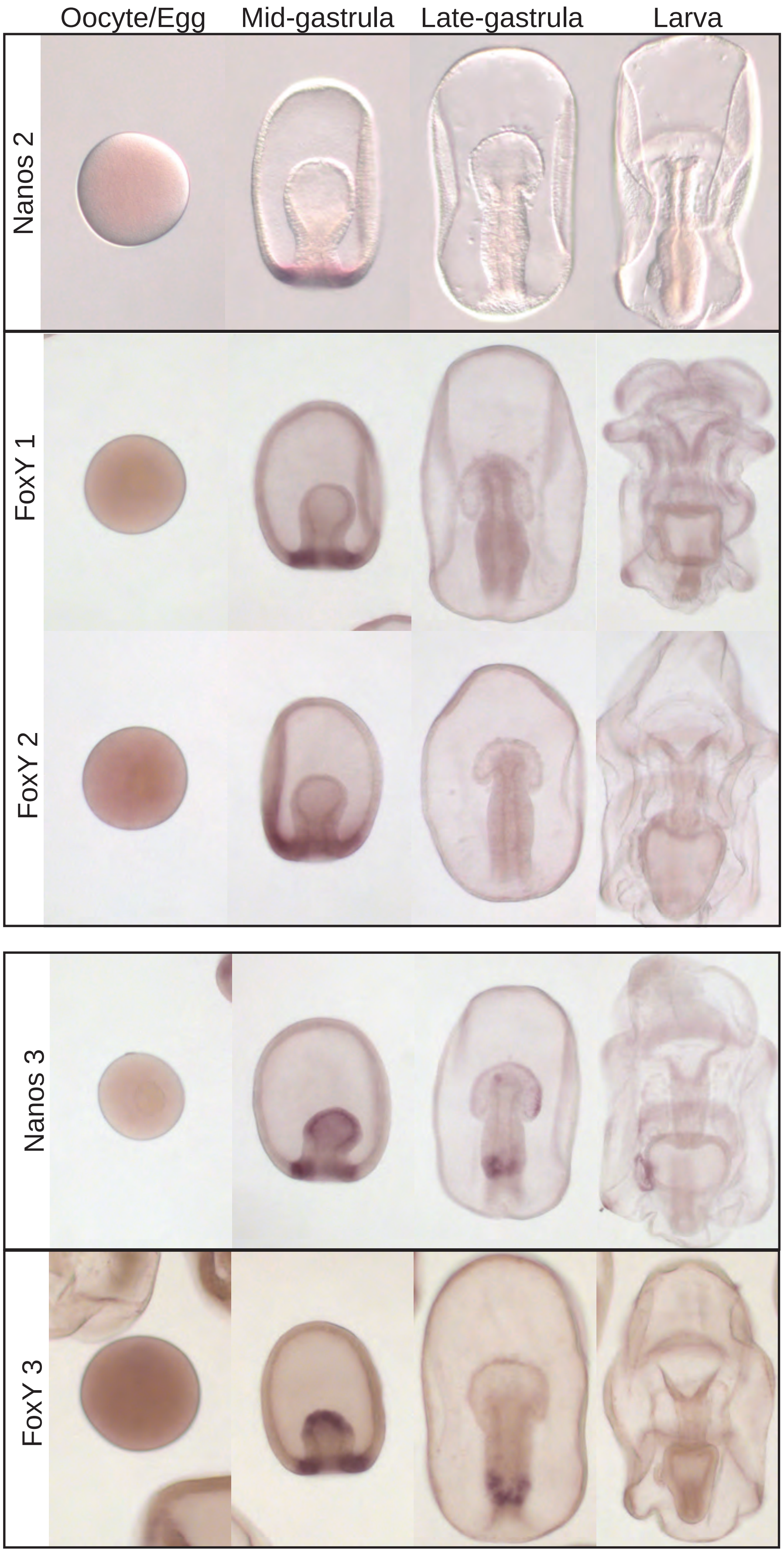
